# Use of F2 bulks in training sets for genomic prediction of combining ability and hybrid performance

**DOI:** 10.1101/501916

**Authors:** Frank Technow

## Abstract

Developing training sets for genomic prediction in hybrid crops requires producing hybrid seed for a large number of entries. In autogamous crop species (e.g., wheat, rice, rapeseed, cotton) this requires elaborate hybridization systems to prevent self-pollination and presents a significant impediment to the implementation of hybrid breeding in general and genomic selection in particular. An alternative to F1 hybrids are bulks of F2 seed from selfed F1 plants (F1:2). Seed production for F1:2 bulks requires no hybridization system because the number of F1 plants needed for producing enough F1:2 seed for multi-environment testing can be generated by hand-pollination. This study evaluated the suitability of F1:2 bulks for use in training sets for genomic prediction of F1 level general combining ability and hybrid performance, under different degrees of divergence between heterotic groups and modes of gene action, using quantitative genetic theory and simulation of a genomic prediction experiment. The simulation, backed by theory, showed that F1:2 training sets are expected to have a lower prediction accuracy relative to F1 training sets, particularly when heterotic groups have strongly diverged. The accuracy penalty, however, was only modest and mostly because of a lower heritability, rather than because of a difference in F1 and F1:2 genetic values. It is concluded that resorting to F1:2 bulks is, in theory at least, a promising approach to remove the significant complication of a hybridization system from the breeding process.

## INTRODUCTION

Since the pioneering work of Shull (1908), hybrid breeding made a significant contribution to increased productivity of globally important field (Duvick 1999) and horticultural (Silva Dias 2010) crops. The development of hybrid varieties rests on the ability to evaluate general combining ability (GCA) of parental inbred lines in earlier stages of the breeding cycle, as well as specific combining ability (SCA), or hybrid performance in general, of particular combinations in stages leading up to release and commercialization (Sprague and Tatum 1942).

Genomic prediction methodology (Meuwissen *et al*. 2001) has successfully been applied to prediction of GCA (Albrecht *et al*. 2011; Würschum *et al*. 2013; Jan *et al*. 2016) and hybrid performance (Massman *et al*. 2013; Technow *et al*. 2014a). This has greatly increased the scale, speed and accuracy of breeding operations (Cooper *et al*. 2014) and promises an increased rate of genetic gain (Heffner *et al*. 2010; Gaynor *et al*. 2017). Building accurate genomic prediction models requires training data sets of hundreds or even thousands of phenotyped and genotyped individuals (Jannink *et al*. 2010; Lorenz 2013; Hickey *et al*. 2014) which can put enormous strain on resources. Promising approaches for increasing the efficiency and throughput of phenotyping (Araus and Cairns 2014; Tardieu *et al*. 2017) and genotyping (Poland and Rife 2012; Gorjanc *et al*. 2016; Technow and Gerke 2017) are currently being developed.

However, producing high-quality F1 testcross seed in sufficient quantities for multi-environment field trials can be an enormous challenge when done for hundreds or even thousands of training individuals. This is the case particularly for autogamous species with hermaphrodite flowers, a group that includes important field crops such as wheat (*Triticum aestivum*), rapeseed (*Brassica napus*), cotton (*Gossypium hirsutum*), sugar beet (*Beta vulgaris*), rice (*Oryza sativa*) and sunflower (*Helianthus annuus*), as well as many horticultural species. Here, hybrid seed production requires elaborate hybridization systems (Kempe and Gils 2011; Whitford *et al*. 2013) that add considerable complexity and cost to the breeding process (Silva Dias 2010; Longin *et al*. 2015). For example, cytoplasmatic male sterility (CMS), one of the most widely used hybridization systems (Janick 1998; Kempe and Gils 2011; Bohra *et al*. 2016; Kim and Zhang 2018), requires (1) development of sterile versions of inbreds (“A-lines”) from the female heterotic pool by backcrossing them into a sterile cytoplasm source, (2) retaining of fertile versions of the same inbreds for seed multiplication (“B-lines”) and (3) introgression of effective fertility restoration genes into the male inbred lines. The complexity and costs arising from applying this and similar systems to hundreds or thousands of individuals for training set development might be prohibitive, particularly when those individuals were chosen with regard to maximum informativeness for model building (Rincent *et al*. 2012) but have no breeding purpose as selection candidates otherwise.

One possible solution for making large scale production of testcross hybrid seed more feasible is to field test bulks of selfed F1 seed (F1:2), instead of the F1 hybrids directly. There are examples of commercial use of F2 hybrid seed in cases where F1 hybrid seed production is not economically feasible (Janick 1998; Wu *et al*. 2004; Bosland 2005). This option has also been considered for application of classical mating designs such as Designs I and II (Comstock and Robinson 1948, 1952) to autogamous species (Stuber 1970). The idea is to test bulked F2 seed from selfed F1 plants obtained by hand pollination (Figure 1). In this system, the seed is produced from vigorous F1 plants instead of inbred lines and according to the natural mode of fertilization (at least for autogamous species). Only very few F1 plants would therefore be required for producing sufficient seed quantities for multi-environment testing and those could easily be produced by hand-pollination, without requiring any form of pollination control such as CMS. Using F1:2 bulks instead of F1 hybrids could therefore be a cost efficient option for producing testcross seed for the hundreds or thousands of individuals comprising training sets in genomic prediction. Under non-additive gene action, however, F1 and F1:2 performance are expected to differ. Using predictions obtained from F1:2 based training sets could then negatively impact genetic gain at the F1 level, which remains the selection target. The objective of this study is therefore to evaluate the prospects of using F1:2 bulks for genomic prediction of F1 hybrid and GCA performance with quantitative genetic theory and stochastic simulation of a genomic prediction experiment.

**Figure 1.**
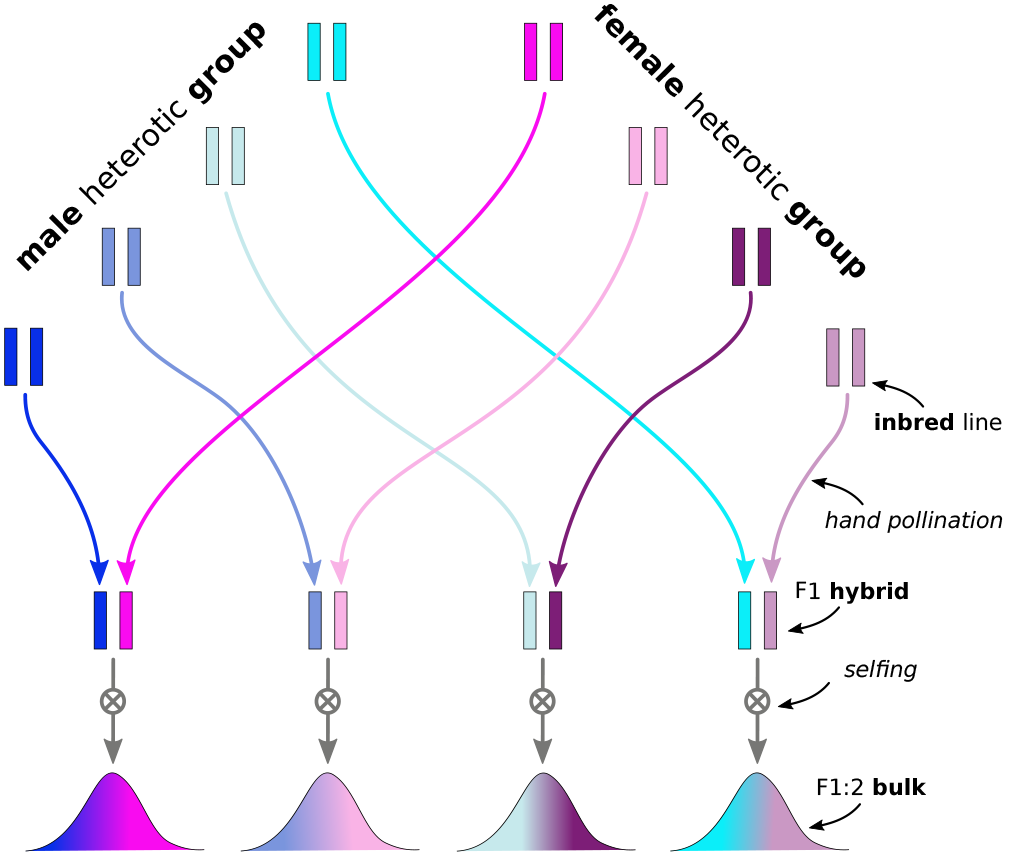
Schematic visualization of creation of F1:2 bulks of F1 interpopulation hybrids.

## MATERIALS AND METHODS

### Theory

Assume a heterotic pattern formed by two populations (“heterotic groups”), arbitrarily labelled “male” (Π_*m*_) and “female” (Π_*f*_). The members of both populations are fully homozygous inbred lines, either produced as doubled haploids (DH), the method of choice in many crop species (Dwivedi *et al*. 2015), or by repeated selfing to a degree that residual heterozygosity is negligible. Consider further two biallelic, independent loci in linkage equilibrium, with alleles *B*_1_ and *B*_2_ and *C*_1_ and *C*_2_, respectively. A superscript *m* or *f* will be used to indicate the origin of the allele (e.g., 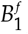 when the allele *B*_1_ originates from Π_*f*_). The alleles are assumed identical in state (i.e., biological function) in both populations (e.g., 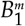 is identical in state with 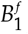). Let 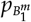 and 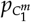 and 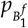 and 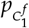 denote the frequencies of the 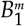 and 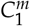 alleles in Π_*m*_ and of the 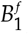 and 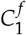 alleles in Π_*f*_, respectively. The frequencies of the alternate alleles are, for example, 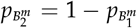. Allele frequencies might differ in Π_*m*_ and Π_*f*_.

When the populations Π_*m*_ and Π_*f*_ are intermated strictly at random, the resulting set of P_*m*_ × Π_*f*_ F1 hybrids forms a “gene-orthogonal population” (Schnell 1965; Melchinger *et al*. 2005). With two biallelic loci and alleles defined according to their origin, there are 16 distinct genotypes, indexed as 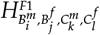. For notational simplicity this will be shortened to 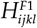. In a gene-orthogonal population the genotype frequencies of 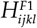 follow from the products of the allele frequencies in Π_*m*_ and Π_*f*_, e.g., the frequency of 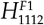 is 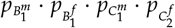. These frequencies will be denoted by *P_ijkl_*.

Selfing each member of 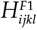 and “bulking” the progeny results in a set of F1:2 bulks denoted by 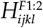 (i.e., the F2 seed from each F1 is bulked separately for each member of 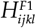. With alleles defined according to origin, each individual 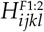 bulk comprises 16 distinct genotypes, each with genotype frequency of 1/16 when assuming absence of segregation distortion. The frequencies of the different 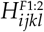 bulks themselves are also *P_ijkl_*.

Consider a three by three matrix ***U*** with elements *u_xy_* equal to the genotypic value of the two-locus genotypes formed by the xth genotype of the ’B’ locus and the yth at the ’C’ locus. The homozygous *B*_1_*B*_1_ genotype thereby corresponds to row *x* = 1, the heterozygous *B*_1_*B*_2_ or *B*_2_*B*_1_ genotype to *x* = 2 and the alternate homozygous to *x* = 3, correspondingly for the *C* locus (see below for examples). Because alleles are assumed to be identical in state in both populations, it is not necessary to distinguish them by origin for the purpose of describing possible genotypic values. Let 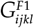 denote the genotypic values of the members of 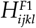. The rows *x* and columns *y* of ***U*** corresponding to elements in 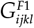 are *x* = *i* + *j* − 1 and column *y* = *k* + *l* − 1.

Similarly, let 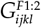 denote the average genotypic values of the F1:2 bulks 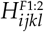. Those can also be obtained from ***U*** as the average genotypic value of the possible genotypes in the bulk, weighted by their frequencies, when assuming absence of segregation distortion. For this purpose, the origin of the allele is again not distinguished. For example, 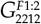 would equal 0.25*u*_31_ + 0.5*u*_32_ + 0.25*u*_33_.

The correlation between 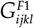 and 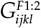 (*cor_hybrids_*), a measure for the similarity between both, was calculated according to the standard statistical definition (e.g., Mood *et al*. 1973), with the mean of 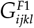 being 
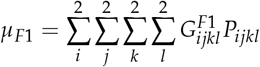
and the variance 
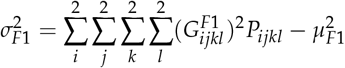
and with *µ_F_*_1:2_ and 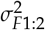 obtained analogously. Thus, 
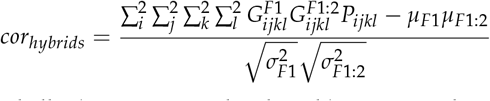

These and all other equations developed here are implemented in a worksheet that is available as supplemental material (File S1).

The GCA of a male inbred with genotype *B_i_C_k_* was evaluated as 
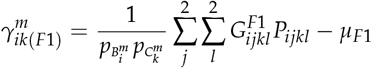
with the corresponding values for the other population and those for the F1:2 bulks (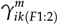) being defined accordingly. The correlation between 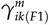 and 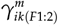 was used to assess the similarity of GCA effects evaluated in F1 hybrids and F1:2 bulks and will be denoted as *cor_GCA_*_(*m*)_ and *cor_GCA_*_(*f*)_ for Π_*m*_ and Π_*f*_, respectively. These correlations were obtained analogously to *cor_hybrids_* and according to their standard statistical definition (e.g., Mood *et al*. 1973). The variances of 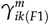 etc., which are required for computing the correlations, will be denoted as 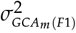 etc. Because populations Π_*m*_ and Π_*f*_ are intermated at random, the male and female effects are uncorrelated with each other. The variance of SCA effects can then be calculated as 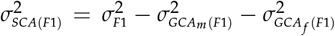, similarly for 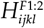.

The quantities 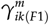, etc., are statistical effects that depend on the allele frequencies in both populations. Thus, two members of Π_*m*_ and Π_*f*_ with identical genotypes will have different GCA values (e.g., 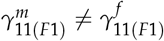), unless both populations have identical allele frequencies, even though the alleles are assumed to be identical in state in both populations (Schnell 1965; Stuber and Cockerham 1966). Therefore also *cor_GCA_*_(*m*)_ ≠ *cor_GCA_*_(*f*)_ in general. For simplicity, however, the average across both, denoted as *cor_GCA_* will be used, where appropriate.

#### Models of genetic architecture and population structure

Several models of gene action will be considered, all encoded through ***U***. The first involves only additive and dominant gene action (“dominance model”), here ***U*** is 
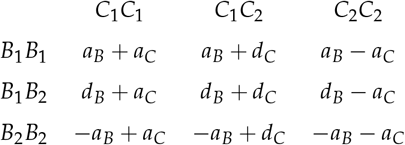
where homozygous effects for the B and C locus, *a_B_* and *a_C_*, and heterozygous effects *d_B_* and *d_C_* are defined according to Falconer and Mackay (1996). A purely “additive model” follows by setting *d_B_* = *d_C_* = 0. Here, however, 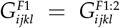 in the absence of segregation distortion and so *cor_hybrids_* and *cor_GCA_* are equal to one. The same is the case for the “additive by additive” epistatic model without dominance or epistatic interactions involving dominance (Hill *et al*. 2008): 
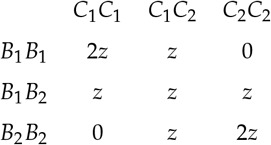
where *z* is an arbitrary constant. The “duplicate factor model” involving additive and dominant gene action as well as all forms of epistatic interactions (Hill *et al*. 2008) is 
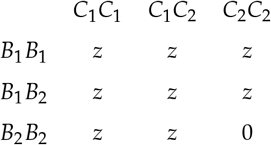
and the “complementary model”, also involving all forms of gene action (Hill *et al*. 2008), is 
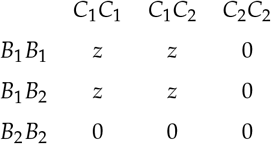

Biological interpretations and examples for the latter two models are given by Holland (2001).

For the duplicate and complementary epistasis models, the quantities 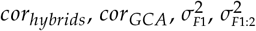 as well as the proportion of SCA to total genetic variance (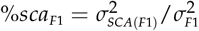 and 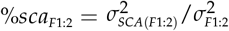) were evaluated across a dense grid of degree of allele frequency differences between Π_*m*_ and Π_*f*_ for the *B* and *C* locus. This difference will henceforth be referred to as “allele divergence” and defined as the difference between, e.g., 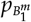 and 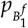, with the midpoint being 0.5. Thus, at a divergence of 0.20, 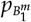 = 0.6 and 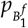 = 0.4, for example. For both models, *z* = 1 was used. Only one locus was considered for the dominance model (i.e., *a_C_* = *d_C_* = 0) and in addition to the allele divergence, the degree of dominance was varied from 0.0 to 3.0 (the homozygous effect was kept constant at *a_B_* = 1).

### Simulation of genomic prediction experiments

A comprehensive simulation of genomic prediction experiments was carried out to evaluate the accuracy of genomic models fitted from training data sets of F1 hybrids and F1:2 bulks for the purpose of predicting hybrid and GCA performance at the F1 level.

#### Parental inbred line genomes

The observed genotypes at 35,478 single nucleotide polymorphism (SNP) markers of 209 maize inbred lines from the Dent (123) and Flint (86) heterotic groups of the maize breeding program of the University of Hohenheim in Germany formed the starting point of the simulation. This data set is available publicly as supplemental material to a publication by Technow *et al*. (2014a). This data was chosen to ensure that the simulated experiments are reflective of the genome properties (allele frequency distribution, linkage pattern and population structure) of an applied hybrid breeding program. The genome properties as well as the history of these populations and the heterotic pattern they form were described in previous studies (Technow *et al*. 2014a,b). For consistency sake, the Dent and Flint populations will arbitrarily be referred to as “male” and “female”, respectively.

#### In-silico biparental populations

Biparental families of DH lines derived from elite inbred parents are the predominant population type encountered in early stages of breeding programs (Mikel and Dudley 2006; Riedelsheimer *et al*. 2013; Hickey *et al*. 2015). In the simulation, each heterotic group was represented by 40 biparental families. The process of how those were created will be described for the male group but was followed analogously for the female group. Five of the 123 male inbred lines were assigned to the “high importance” group, twenty to the “medium importance” group and the remaining 98 to the “low importance” group. This assignment was done at random. The “high importance” inbreds were given a weight of 0.1, the “medium importance” inbreds a weight of 0.0125 and the remaining inbreds a weight of 0.00255 (the weights of all inbreds sum to 1.0). Then all 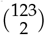 possible biparental crosses among those 123 inbreds were assigned weights equal to the product of the weights of the corresponding parents. The 40 biparental families were then drawn at random from all possible families with probabilities proportional to the weights previously assigned. This results in an unbalanced contribution of inbred lines to the breeding populations and reflects that often just a few, very successful inbred lines are used disproportionately by breeders (Mikel and Dudley 2006; Technow *et al*. 2014b).

Then, 25 recombinant DH lines were generated in-silico from each family by simulating meiosis between the gametes of the parents followed by a chromosome doubling step. Meiosis was simulated according to the Haldane mapping function with the software package “hypred” (Technow 2013). Thus, each heterotic group comprised 1,000 DH lines from 40 biparental families. A random sample of 15,000 of the SNP markers was used for these simulations to facilitate computations. Because simulation of meiosis requires a genetic linkage map, the physical map positions of the SNP loci where rescaled linearly to the chromosome lengths of the genetic map reported by Fu *et al*. (2006).

#### Simulation of genetic architecture

200 loci were defined to be QTL with direct influence on a generic complex trait. Those loci were chosen from the set of 15,000 SNP included in the simulation according to two population structure scenarios: “convergent” and “divergent”. In the convergent scenario, the QTL had similar allele frequencies in both heterotic groups with a maximum absolute difference of 0.05 (i.e., |*p_m_* − *p_f_*| < 0.05). This corresponds to a newly formed hybrid breeding program before the establishment of distinct heterotic groups (Melchinger 1999; Fischer *et al*. 2009). In the divergent scenario, both populations had very different allele frequencies with a minimum absolute difference of 0.60 (i.e., |*p_m_* − *p_f_*| > 0.60), which corresponds to a well established hybrid breeding program in which heterotic groups have diverged as a result of many cycles of reciprocal recurrent selection (Labate *et al*. 1999; Reif *et al*. 2007; Technow *et al*. 2014a; Larièpe *et al*. 2017). The definitions of the convergent and divergent heterotic group scenarios correspond to those in Technow *et al*. (2012). An additional requirement was that loci used as QTL had to have a minimum minor allele frequency of 0.025 in each heterotic group to ensure that they were contributing to genetic variation. Within those constrains, the 200 QTL were drawn at random.

They were then randomly separated into 100 two-loci pairs. Each pair was assigned a matrix ***U*** describing the genotypic values. On average, 5% of the loci were assigned to the additive and another 5% to the additive by additive gene action models. The dominance model was assigned to 10% of the pairs. The remaining 80% of pairs were assigned to the complementary and the duplicate factor gene action models in equal proportion. Thus, on average 90% of the QTL gave rise to non-additive gene action effects that affect F1 and F1:2 genetic values differently. Note, however, that the latter three gene action models contain all types of gene action effects, including additive and additive by additive.

The homozygous effects *a* (used for the additive and dominance models) were drawn from a Normal distribution with mean zero and standard deviation of 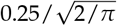 (throughout, the Normal distribution will be parametrized by its mean and standard deviation). Their absolute values then have an expectation of 0.25. The heterozygosity gene effects *d* were drawn from 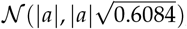, resulting in an average degree of dominance of one, which is consistent with experimental results in hybrid crops (Gardner 1963; Radoev *et al*. 2008; Schön *et al*. 2010; Larièpe *et al*. 2012). Using 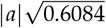 as standard deviation ensures that 90% of the sampled heterozygosity effects are above zero. This was done because dominance effects for traits showing hybrid vigour tend to be positive (Semel *et al*. 2006; Bennewitz and Meuwissen 2010; Huang *et al*. 2015). For the additive by additive model, the value of *z* was drawn from 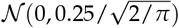 and for the duplicate and complementary gene action models from 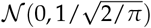. The absolute values of *z* thus have expectations of 0.25 and 1.00, respectively. Those settings for the distributions of *a, d*, and *z* were chosen to ensure that the various gene action effects have the same magnitude in all gene action models. For example, with *z* = 1, *a* = *d* = 0.25 in the duplicate and complementary gene action models and hence equal to their expected values in the additive and dominance gene action models. Gene action effects for arbitrary ***U*** can be calculated with File S1 according to definitions by Holland (2001). Because interaction systems among loci can rarely be cleanly assigned to a particular model (Holland 2001), a small amount of “genetic noise” was added to the elements of each ***U*** matrix. Those values were drawn from a Normal distribution with mean zero and standard deviation equal to 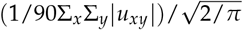. The mean absolute value of those deviations was thus expected to be equal to 1/10th of the mean absolute value of the elements of a particular instance of ***U***. All ***U*** matrices thus slightly deviated from their assigned models of gene action and as a result, even pairs assigned the simple additive model will give rise to small amounts of variation due to dominance and epistatic gene action.

#### In-silico population of hybrids

The genotypic values of all F1 hybrids from the full factorial of 1, 000 × 1, 000 = 1, 000, 000 interpopulation hybrids were calculated by summing the genetic effects across all 100 two-loci pairs according to the QTL genotypes of the hybrids. Because the parents were fully homozygous DH lines, the QTL genotypes of the hybrids follow directly from the genotypes of their parents. This full factorial was defined as the reference population of hybrids. The true GCA values of all 1,000 male and 1,000 female DH lines were calculated from the row and column means of the full factorial table (Sprague and Tatum 1942). Accordingly, the true SCA effects of all hybrids were obtained as the difference between the performance of the individual hybrids and the parental GCA effects.

#### Genomic prediction training set

The DH from 20 male and 20 female families, chosen at random from the 40 male and female families created, were used for building the training set. Each of the 500 DH from the 20 families from one heterotic group was paired at random with one DH from the 20 families of the opposite heterotic group, resulting in a set of 500 hybrid combinations (Figure 2). This design is similar to the reciprocal testcrossing design used by Kadam *et al*. (2016). The 500 male and 500 female DH lines as well as the 500 hybrid combinations among them will henceforth also be referred to as “tested”. The 500 hybrid combinations comprising the training set are a subset of the total population of hybrids and the genotypic performance values of the F1 versions were calculated as described before. Phenotypic values were obtained by adding a Gaussian noise term with standard deviation chosen in such a way that the broad sense heritability was 0.5. This training set will also be refered to as F1 training set. The alternative F1:2 training set was made up of F1:2 bulks created from the same hybrids that were used for the F1 training set. For this, each F1 was selfed in-silico and 100 F2 individuals generated by simulating meiosis as described before. The true genetic values of each of the 100 F2s was calculated similarly as for the F1s and then averaged to arrive at the genetic performance of the F1:2 bulk. Phenotypic values were generated according two scenarios. In the first, the residual variation used for the F1 training set was assumed constant (“constant residual variation” scenario), in the second a constant heritability of 0.5 was assumed also for the F1:2 estimation set by adjusting the residual variation accordingly (“constant heritability” scenario).

**Figure 2.**
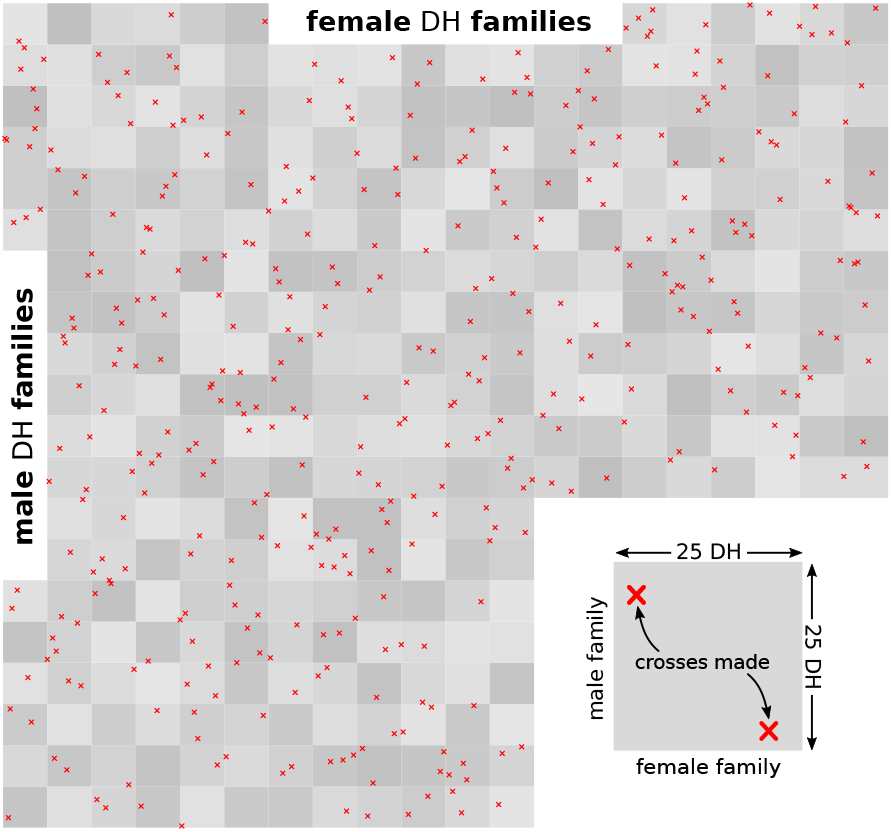
Schematic visualization of reciprocal crossing design used to build the training set.

#### Prediction model

The following mixed model was fitted to the data 
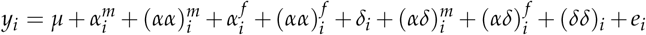
where *y_i_* is the scaled and centred phenotypic value of the *i^th^* training set entry (either a F1 hybrid or a F1:2 bulk, depending on the training set). The intercept is denoted by *µ*. Male and female additive main and additive by additive epistatic interaction effects are 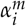 and 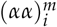, and 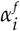 and 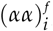, respectively, which together constitute the GCA of a fully homozygous male and female parents of the *i^th^* hybrid combination (Falconer and Mackay 1996, p. 276). Dominance effects are denoted by *d_i_*, interaction effects between male and female additive and dominance effects are 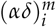 and 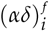, respectively, and (*δδ*)_*i*_ denote the dominance by dominance interaction effects. Together, those effects constitute the SCA effect of the *i^th^* hybrid combination. The residual associated with *y_i_* is *e_i_* and was modelled as iid 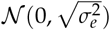. The other effects were modelled by Multivariate-Normal distributions with mean vectors of zero and covariance matrices defined as 
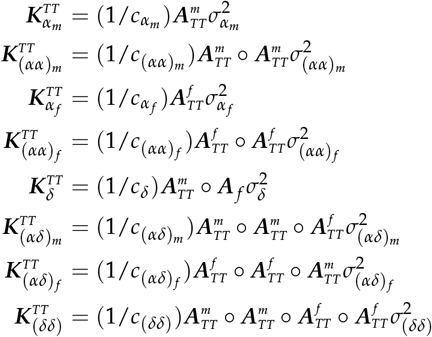
following Stuber and Cockerham (1966), where 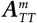 and 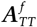 are the additive genomic relationship matrices of the tested male and female parents, respectively, of the training set hybrids and ’○’ indicates element-wise multiplication. These matrices were calculated from the marker data following VanRaden (2008) as 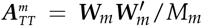, where *M_m_* is the number of markers and 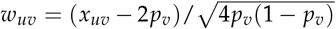 (with *u* indexing the parent and *v* the marker), *x_uv_* coding the number of reference alleles (taking values of 0 or 2), and *p_v_* being the allele frequency of the reference allele in the male population. The genomic relationship matrix of the female parents was obtained analogously. Both matrices were calculated from the same set of 5,000 marker loci, which were obtained by randomly sampling from the initial set of 15,000 loci but excluding the 200 loci defined as QTL. The terms *c_α_m__* etc. are normalization factors equal to 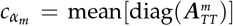, etc. and help to bring the estimated variance components onto a comparable scale with the residual variance (Xu 2013). This model is similar to the one used by Massman *et al*. (2013) and Technow *et al*. (2014a) except that these authors did not consider epistatic effects. The model was fitted using the R package ‘BGLR’ (Pérez and Campos 2014) and its default settings for prior distributions and hyperparameters. A total of 1,000 samples were obtained from a chain of length 100,000, with a burn-in of 50,000 and a thinning interval of 50. The posterior means of the variance components and of the intercept were used as point estimates.

#### Genomic prediction accuracy

The performance of all F1 hybrids from the full factorial was predicted using BLUP as 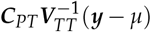 following Henderson (1973), where ***C**_PT_* is the genetic covariance matrix of predicted and tested hybrid combinations and ***V**_TT_* the phenotypic covariance matrix of the data. The elements of ***C**_PT_* and ***V**_TT_* were computed according to Bernardo (1996), with the addition of the covariance matrices of epistatic effects not considered there. Specifically, 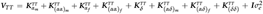. Further, 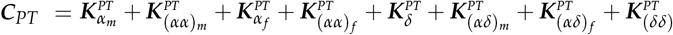 with 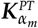 etc., calculated analogously to the corresponding 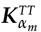 etc. but using additive relationship matrices 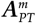 and 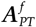, which represent the additive relationships of the parents of the 1,000,000 hybrid combination to be predicted and the parents of the training set hybrids. The normalization constants calculated previously were used to normalize these matrices, too. Predictions for SCA effects were obtained by limiting ***C**_PT_* to the covariance matrices of the effects contributing to SCA. Following Technow *et al*. (2012), hybrids for which both the male and the female parent were tested were assigned to the “T2” prediction set, hybrids for which either the male or the female parent were tested (but not both) were assigned to the “T1” set and hybrids without any tested parent were assigned to the “T0” set. The tested hybrids themselves were assigned to the “T3” set. Prediction accuracy of total hybrid performance and SCA effects was defined as the Pearson correlation coefficient between predicted values and the corresponding true values of the F1 hybrids. To emphasize, the true performance and SCA effects of the F1 hybrids were used as reference values also when assessing the accuracy of the model fitted from the F1:2 training set.

When predicting the GCA of DH lines, 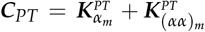 for males and 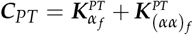 for females. The matrices 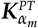 etc., were computed as before but from 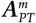 and 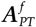 which for this purpose contained the relationships between all 1,000 male or female parents and the male or female parents of the estimation set hybrids. Prediction accuracy of GCA effects was defined as the Pearson correlation coefficient between predicted and true GCA effects obtained from the full factorial of F1 hybrids. Thus, the prediction accuracy evaluates the ability to predict F1 based GCA, even when the F1:2 training set is used. The prediction accuracy was calculated separately for tested and untested DH lines and within and across families. The within family accuracies were averaged across families. To simplify the results further, all accuracies were averaged across male and female heterotic groups.

The whole simulation was repeated independently 2,500 times for each heritability and population structure scenario and results averaged. All computations were conducted within the R statistical computing environment (R Core Team 2018).

### Data Availability

The SNP genotypes of the Dent and Flint inbred lines used as the initial population and the physical map of the SNP are available from the online supplement of Technow *et al*. (2014a).

## RESULTS

### Theoretical models

The correlation between F1 hybrids and F1:2 bulks (*cor_hybrids_*) was mostly high (above 0.9) in the dominance (Figure 3A) and both epistatic models (Figures 4A and 4G). The widest range of values was thereby observed for the duplicate model, where *cor_hybrids_* reached below 0.5 when allele divergence was extreme (Figure 4A). In the dominance model *cor_hybrids_* decreased with increasing degree of dominance, but remained high throughout. In the complementary model it was lowest when divergence was high at both loci, but again only slightly below the highest values, which were achieved when none or only one of the loci were strongly diverged (Figure 4G). The correlation between GCA effects obtained from F1 or F1:2 (*cor_GCA_*, average across male and female correlations) in the duplicate model followed a similar trend as *cor_hybrids_* (Figure 4B) but remained considerably higher also under extreme divergence. In the complementary model *cor_GCA_* was high and confined to a narrow range, similar to *cor_hybrids_* (Figure 4H). The highest values were observed when divergence was either similar at both loci or very different. However, minimum and maximum values were only marginally different. With just a single locus, *cor_GCA_* in the dominance model is either 1.0 or 0.0, depending on the combination of allele divergence and degree of dominance (Figure 3B). More specifically, *cor_GCA_* = 0.0 is a result of the correlation being 1.0 in the population with higher allele frequency and −1.0 in the other. Values of −1.0 are thereby observed within a narrow band defined by particular combinations of allele divergence and degree of dominance. The genetic variance of F1 hybrids 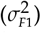 in the dominance model increased with increasing degree of dominance and decreasing divergence (Figure 3C). In the epistatic models, it decreased with increasing divergence at one or both loci (Figures 4C and 4I). Under the dominance and complementary models, the ratio 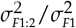 was below one throughout, indicating that F1 hybrids are expected to have a greater genetic variance than F1:2 bulks (Figures 3D and 4J). In the duplicate model the opposite was the case, and 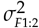 could be considerably larger than 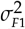 when allele divergence became extreme (Figure 4D). The proportion of SCA to total genetic variance in the F1 hybrids (%*sca_F_*_1_) followed similar trends as the total genetic variance under all models, i.e., it decreased with increasing allele divergence and decreasing degree of dominance (Figures 3E, 4E and 4K). Finally was the ratio %*sca_F_*_1:2_/%*sca_F_*_1_ below one throughout for all three models, indicating that relatively less genetic variation can be attributed to SCA effects in F1:2 bulks than in F1 hybrids (Figures 3G, 4G and 4L).

**Figure 3.**
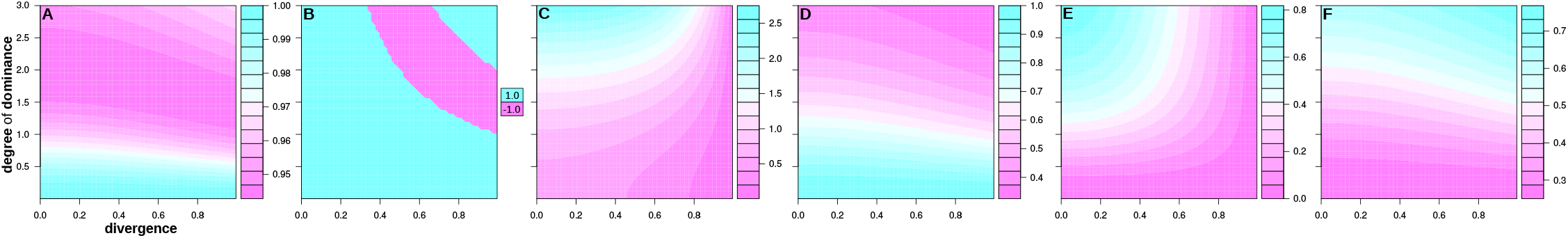
Expected values of *cor_hybrids_* (A), *cor_GCA_*_(*f*)_ (B),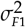 (C), 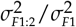 (D), %*sca_F_*_1_ (E) and %*sca_F_*_1:2_/%*sca_F_*_1_ (F) under the “dominance” model as a function of allele frequency divergence and degree of dominance. Note that the colour scale is not constant across sub-figures.

**Figure 4.**
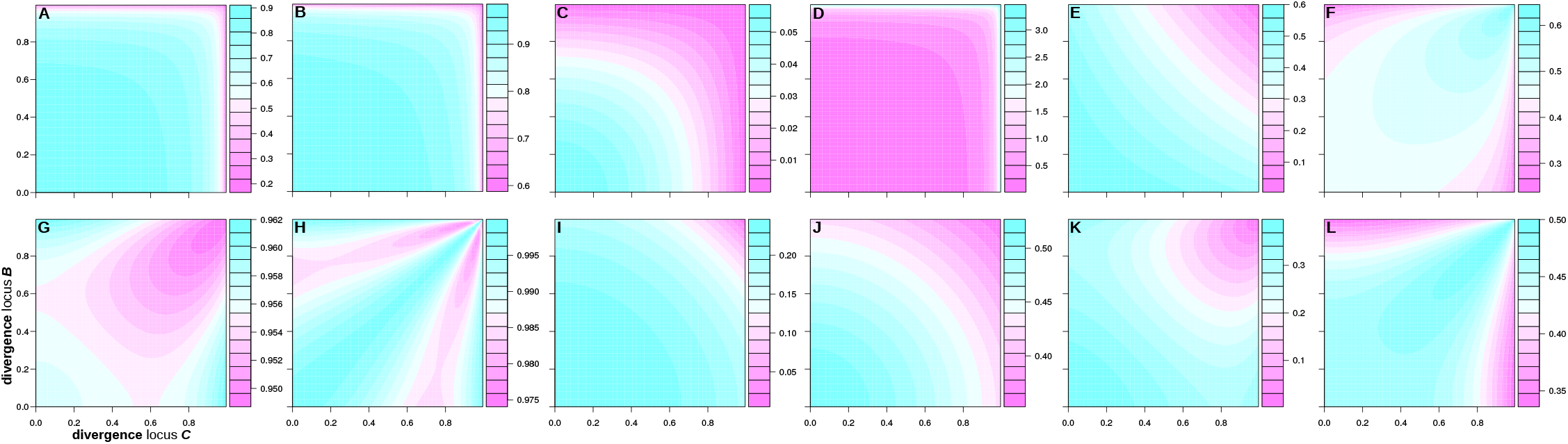
Expected values of *cor_hybrids_* (A,G), *cor_GCA_* (B,H), 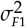 (C,I), 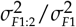 (D,J), and %*sca_F_*_1_ (E,K) and %*sca_F_*_1:2_/%*sca_F_*_1_ (F,L) under the duplicate model (top row) and complementary model (bottom row) as a function of allele frequency divergence at both loci. Note that the colour scale is not constant across sub-figures. The values in sub-figure D were transformed to *ln* 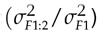 to aid visibility.

### Simulation results

#### Quantitative genetic parameters

Because these parameters are not affected by the heritability scenario, only results for the “constant residual variation” scenario will be shown. The F1 hybrids had a larger genetic variance than F1:2 bulks (Table 1). The difference between the two was relatively larger under the divergent heterotic group structure, where 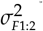 was less than half of 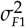. Overall, genetic variance was larger in the convergent structure than in the divergent structure, for both F1 and F1:2. The proportion of SCA to total genetic variation was larger in the convergent than the divergent structure by almost 10 percentage points (Table 1). This quantity could only be assessed for the F1 hybrids for which the full factorial was created. Finally, the correlation between F1 hybrids and F1:2 bulks was largest in the convergent heterotic group structure and very close to 1.00 (Table 1). However, this correlation was with 0.91 very high in the divergent scenario, too.

**Table 1.** Average simulation results of total genetic variance, proportion of SCA variance and correlation between F1 hybrids and F1:2 bulks.

|  | convergent |  | divergent |  |
| --- | --- | --- | --- | --- |
|  | F1 | F1:2 | F1 | F1:2 |
| variance<sup>a</sup> | 20.34 | 12.36 | 15.89 | 7.63 |
| %SCA<sup>b</sup> | 0.24 | — | 0.15 | — |
| correlation<sup>c</sup> | 0.97 |  | 0.91 |  |

<sup>a</sup> True genetic variance of F1 hybrids ( $\sigma_{F1}^2$ ) and F1:2 bulks ( $\sigma_{F1.2}^2$ ) used in the training set
<sup>b</sup> True proportion of SCA to total genetic variance ( $\%sca_{F1}$ ) in full factorial (only measured for F1 hybrids for which full factorial was created)
<sup>c</sup> Correlation ( $cor_{hybrids}$ ) of true genetic values of F1 hybrids and F1:2 bulks in training set

#### GCA prediction

Under constant residual variation, the prediction accuracy obtained from the F1 training set was higher than that for the F1:2 training set throughout (Table 2). Under a convergent heterotic group structure, the differences were small and did not exceed 0.04 points. Differences under the divergent structure were in the magnitude of 0.10 points. When the heritability of the F1:2 bulk phenotypes was made equal to that of the F1 hybrids (constant heritability scenario), the GCA prediction accuracy from the F1:2 training set increased considerably and now was close to (divergent structure) or even slightly higher (convergent structure) than that of the F1 training set. (The heritability of the latter was 0.5 in both cases and so the accuracy values were not expected to change.) Overall, prediction accuracy was higher for DH lines that contributed to the training set (“tested”) than for DH lines that did not (“untested”) and lower within families than across.

**Table 2.** Average simulation results for genomic prediction accuracy of GCA effects.

| structure | generation | within family |  | across families |  |
| --- | --- | --- | --- | --- | --- |
|  |  | tested | untested | tested | untested |
| — **constant residual variation<sup>a</sup>** — |  |  |  |  |  |
| convergent | F1 | 0.49 | 0.34 | 0.71 | 0.51 |
|  | F1:2 | 0.45 | 0.32 | 0.67 | 0.49 |
| divergent | F1 | 0.50 | 0.36 | 0.73 | 0.54 |
|  | F1:2 | 0.40 | 0.29 | 0.62 | 0.46 |
| —— **constant heritability<sup>b</sup>** —— |  |  |  |  |  |
| convergent | F1 | 0.49 | 0.34 | 0.71 | 0.51 |
|  | F1:2 | 0.51 | 0.36 | 0.72 | 0.52 |
| divergent | F1 | 0.50 | 0.36 | 0.73 | 0.54 |
|  | F1:2 | 0.48 | 0.34 | 0.69 | 0.51 |

<sup>a</sup> Phenotypic values in F1 and F1:2 training sets had same magnitude of residual variation
<sup>b</sup> Phenotypic values in F1 and F1:2 training sets had same heritability

#### Hybrid prediction

Under constant residual variation, the prediction of hybrid performance was more accurate when using the F1 training set than for the F1:2 training set (Table 3). The difference between the two thereby was largest for the divergent heterotic group structure, where it reached from 0.15 points for T3 hybrids to 0.06 points for T0 hybrids. Under the convergent heterotic group structure, the differences reached from 0.09 points (T3) to 0.01 points (T0). With constant heritability the accuracy of the predictions from the F1:2 training set increased markedly and now were similar (divergent structure) or even slightly higher (convergent structure) than for the F1 training set. In general, the performance of T3 hybrids was predicted with highest accuracy, followed by T2, T1 and T0 hybrids. The prediction accuracy of SCA effects was considerably lower than that of total hybrid performance throughout (Table 3). Similar trends held, however, with the exception that even under constant heritability the F1:2 training set was considerably less accurate than the F1 training set.

**Table 3.**
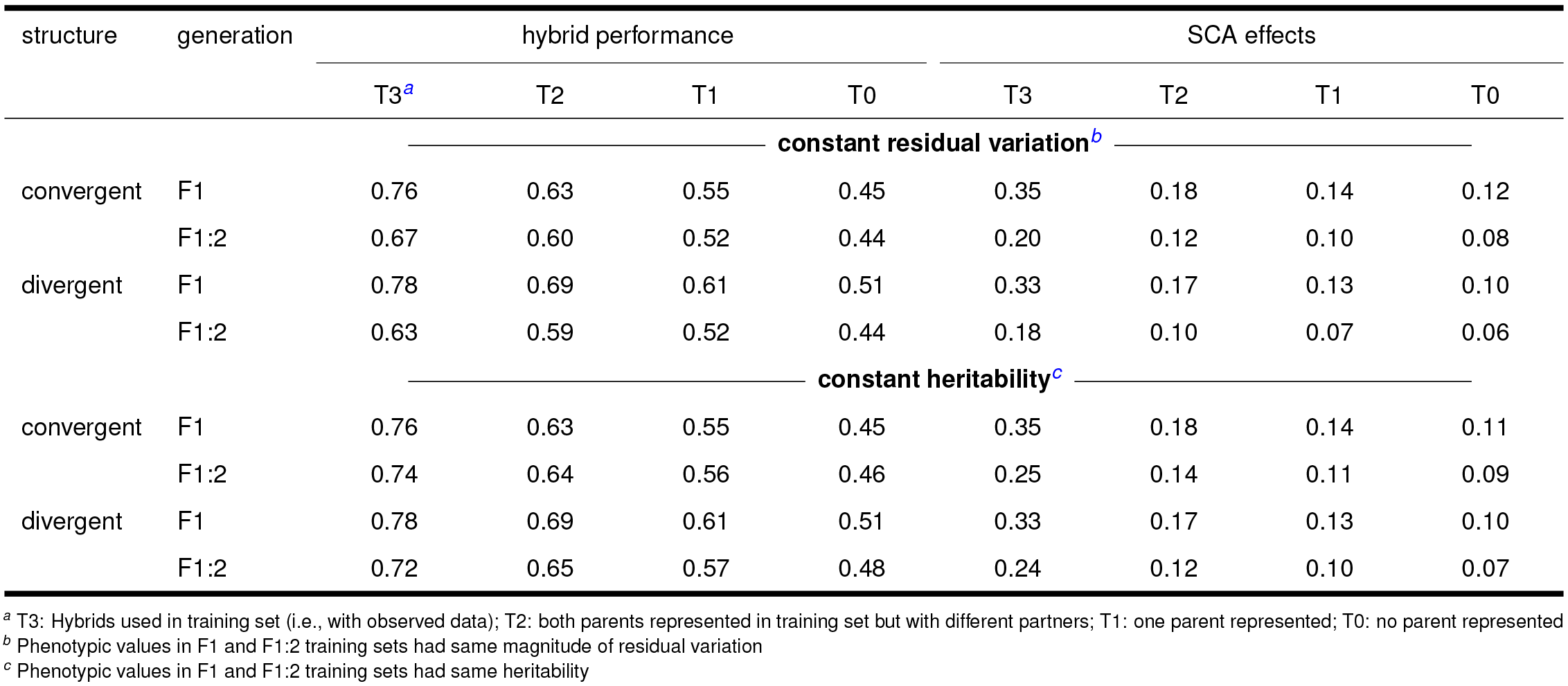
Average simulation results for genomic prediction accuracy of hybrid performance and SCA effects.

## DISCUSSION

Using F1:2 bulks for genomic prediction training sets and more generally testcross and hybrid evaluation is an alternative when production of large quantities of seed of F1 hybrids is expensive, significantly increases the complexity of a breeding program or is impossible altogether. The objective of this study was to assess the promise of this approach from a theoretical point of view with the help of quantitative genetics theory and stochastic simulation.

### Quantitative genetic properties

Both the simulation as well as the theoretical results indicate that the correlation between F1 hybrids and F1:2 bulks can be expected to be high across a wide range of scenarios (Table 1, Figures 3 and 4). A few exceptions should be pointed out, however. Lower values, particularly for *cor_hybrids_*, were observed for increasing divergence between the allele frequencies in the two heterotic groups, particularly for the duplicate epistatic model (Figure 4A). Here, most of the F1 hybrids are expected to have genotypic value of *z*, and only a small fraction a value of zero (those homozygous for the ’2’ allele at both loci). F1:2 bulks derived from F1 hybrids heterozygous at one or both loci, however, will exhibit an average performance that is intermediate between *z* and 0.0. With increasing allele frequency divergence, relatively more F1 hybrids are heterozygous at one or both loci, leading to a greater frequency of cases with differing F1 and F1:2 performance and hence a lower correlation.

Two factors will lesson the impact of strongly diverged pairs of loci with duplicate gene action on the overall correlation for complex traits. Firstly, is the genetic variance generated by loci pairs with duplicate gene action considerable lower than for pairs of loci with complementary gene action but similar divergence and magnitude of gene action effects (compare figures 4C and 4I). Secondly, does the generated variance decrease with increasing divergence (Figure 4C). Thus, when a trait is influenced by both duplicate and complementary epistasis and by loci with different degrees of divergence, the impact of strongly diverged loci exhibiting duplicate epistatic on the overall correlation between F1 and F1:2 genetic effects will be relatively minor.

Under dominant gene action, the correlation between F1 and F1:2 GCA effects in the heterotic group with lower frequency of the allele that increase genetic value can take values of –1 within a narrow band defined by particular combinations of allele divergence and degree of dominance (Figure 3B). An explanation for this phenomenon is provided in the supplemental file S2. For brevity it should suffice here to state that for degrees of dominance below one (i.e., partial dominance), the correlation is always +1 and that very high degrees of dominance are required to reach this “band” for low or moderate divergence of allele frequencies. As can be seen from the example in File S2 (and from File S1) as well, loci that exhibit this behaviour will contribute relatively little to the variance of GCA effects in the affected heterotic group. Thus, to have an impact on the overall GCA correlation, the vast majority of loci controlling a complex trait must be within the specific combinations of allele divergence and degree of dominance defining this “band” and the alleles increasing genotypic value must consistently have a lower frequency in the same heterotic group. This seems unlikely for a complex trait controlled by hundreds or thousands of genes. The low contribution to GCA variance in the heterotic group in which *cor_GCA_* is –1 also explains why *cor_hybrids_* remains high even then.

The theoretical results show that to the degree that *cor_hybrids_* (and *cor_GCA_*) did decrease, it largely did so as a function of increasing allele divergence. This explains that *cor_hybrids_* was lower in the divergent heterotic group structure, characterized by strong differences in male and female frequencies of QTL alleles, than in the convergent structure, where QTL alleles were constrained to have similar frequency in both heterotic groups.

The observation that the genetic variance is lower among F1:2 bulks than among F1 hybrids (Table 1) is also in line with the theory, which further predicts that this difference increases as allele frequencies diverge. This again is observed in the simulation results, in which the F1:2 had approximately 61% of the variation of the F1 in the convergent heterotic group structure but only about 48% in the divergent structure. Duplicate epistasis again presents somewhat of an exception, because here the F1:2 actually had a larger variance than the F1 (Figure 4D) and the more so the more the alleles diverged. As mentioned before, however, the weight of loci with duplicate epistatic effects on determining overall genetic trends for complex traits is considerably lower than that of loci with other types of gene action.

Finally, the theoretical results show that the proportion of SCA to total genetic variation in the F1 hybrids decreases with increasing interpopulation divergence for the dominance (Figure 3E), the duplicate (Figure 4E) and the complementary model (Figure 4K). This was also observed in the simulation, where the relative amount of SCA variation was almost 10 percentage points higher in the convergent than in the divergent population structure scenario. Reif *et al*. (2007) previously showed that increasing interpopulation divergence reduces the proportion of SCA variance when considering only dominance. For random mating populations in Hardy-Weinberg equilibrium it was further shown that genetic variance generated by the epistatic and dominance models considered here is largely additive when allele frequencies are at extreme values (Hill *et al*. 2008). Theoretical results also imply that the contribution of SCA to total variance is expected to be relatively lower for F1:2 bulks than F1 hybrids. This is easiest to see under dominant gene action, where *d* is in essence halved because only 50% of the segregants within a F1:2 bulk derived from a heterozygous F1 will have a genetic value of *d* and the genetic values of the homozygous segregants will cancel in the determination of the mean value of the bulk.

### Genomic prediction of GCA effects

Based on the developed theory, F1:2 bulks are expected to have a lower genetic variance than F1 hybrids. With constant residual variation, i.e., F1 and F1:2 training sets phenotyped with same number of locations and replications, the latter is thus expected to have a lower heritability. Prediction accuracy is expected to decrease with decreasing heritability of the training set phenotypes (Daetwyler *et al*. 2010; Lorenz 2013). The reduced prediction accuracy coming from the F1:2 training set in the “constant residual variation” scenario (Table 2) thus was in part due to the lower heritability of the F1:2 phenotypes. Indeed, when both training sets had the same heritability, accuracy was similar, too. In the convergent heterotic group scenario the accuracy of the F1:2 training set was then even slightly higher than that of the F1 training set. SCA effects essentially act as noise and even modelling them is unlikely to completely remove their confounding effects on the estimation of GCA effects. This surprising result can therefore be explained by the relatively lower amount of SCA variation in the F1:2 training set, as indicated by the theoretical results (Figures 3F, 4F, and 4L). SCA variation is expected to be relatively less important under a divergent heterotic group structure, as shown by theory and simulation results obtained here and elsewhere (Reif *et al*. 2007) as well as in practice (Fischer *et al*. 2009). The relatively lower amount of SCA variation in the F1:2 training set will therefore be less of a benefit in this case.

Even if the genomic model fitted could explain 100% of the genetic variation in the F1:2 bulks, the prediction accuracy would have an upper bound proportional to the correlation between F1 and F1:2 genetic effects. It was shown here that this correlation is expected to decrease with increasing allele divergence between heterotic groups and observed to be lower lower in the divergent heterotic groups structure than in the convergent structure. Finally, both theory and the simulation show that the relative reduction in genetic variance is greater, the greater the degree of divergence. With a constant residual variation, the reduction in heritability was therefore larger in the divergent scenario. These three factors combined, i.e., the lower benefit from the reduced proportion of SCA variance, the lower correlation between F1 and F1:2 genetic effects, and the lower heritability can explain why a larger accuracy penalty from using F1:2 training sets was observed for the divergent heterotic group structure.

The described trends held in general for prediction of tested and untested DHs. The differences between the accuracy from F1 and F1:2 training sets, however, were larger when predicting tested DH lines than for untested ones (Table 2), at least for the “constant residual variation” scenario. Genomic predictions of tested individuals (i.e., individuals contributing to the training set), are strongly influenced by the phenotypic observations available for them (Endelman and Jannink 2012; Müller *et al*. 2015). It can be speculated that they are therefore more sensitive to differences in F1 and F1:2 genetic values and the lower heritability of the latter. Nonetheless, accuracy for tested DH was always considerably higher than for untested DH. Each tested DH was not only represented in the training set directly in the form of a hybrid progeny, but also had 24 full-sib DH that directly contributed to the training set as well. Previous studies demonstrated the positive effect on prediction accuracy of direct observations (e.g., Endelman *et al*. 2014) and presence of full-sibs in the training set (Habier *et al*. 2013; Riedelsheimer *et al*. 2013; Schopp *et al*. 2017). The trends concerning accuracy of F1 and F1:2 training sets also held for within and across family prediction. That the latter was considerably more accurate was expected because it is largely driven by separation of family means (Windhausen *et al*. 2013; Riedelsheimer *et al*. 2013).

### Genomic prediction of hybrid performance

Because GCA is an important component of total hybrid performance, similar trends were observed here. Namely that the prediction accuracy from the F1:2 training set was trailing that of the F1 training set under constant residual variation but caught up to or even exceeded it under constant heritability (Table 3). Similarly, the accuracy difference was larger for the divergent population structure and for hybrids that were part of the training set (T3). The explanations given above for GCA apply also here. That prediction for T3 hybrids was most accurate, followed by that of T2, T1 and T0 hybrids, was observed in previous studies (e.g., Technow *et al*. 2014a; Zhao *et al*. 2015) and is a result of the decreasing degree of relatedness between the hybrids in the four classes and the training set. SCA represents higher order statistical effects and is therefore estimated with greater error than the GCA effects contributing to hybrid performance (Technow *et al*. 2012; Kadam *et al*. 2016). Their accuracy was therefore considerably lower than that of GCA effects (compare the “across population” section of Table 2 with Table 3). Because it is relatively less important with divergent heterotic groups, the lower accuracy of predicted SCA effect will matter less for determining total hybrid performance, which consequently was predicted with greater accuracy than in the convergent scenario.

There has recently been interest in revisiting reciprocal full-sib mating designs (Hallauer and Eberhart 1970) in light of advances in genomic prediction of hybrid (Technow *et al*. 2012; Kadam *et al*. 2016; Fritsche-Neto *et al*. 2018) and GCA (Giraud *et al*. 2017) performance. As noted by Giraud *et al*. (2017), reciprocal testing has several advantages over the still predominant topcross design, in which individuals under evaluation are crossed with a common partner (“tester”) from the opposite heterotic group (Jenkins and Brunson 1932). A major advantage is that field testing is twice as efficient because testcross hybrids are informative for both heterotic groups, meaning that the same number of individuals can be tested with half the resources (e.g., 1,000 instead of 500 total hybrids would have been required to represent 500 male and 500 female DH if a topcross design would have been used). The other advantage is that when the crosses are made at random, they comply with the assumptions of a gene-orthogonal population (Schnell 1965) thus facilitating unbiased estimation of random GCA and SCA effects and their variances. The training set used in this study was constructed in this reciprocal fashion by directly pairing male and female DH at random, without the use of a topcross tester. The results are thus additional confirmation that hybrid performance as well as GCA and SCA effects can be evaluated and predicted accurately with such a design, even when each parent is represented in only a single hybrid combination.

### Practical considerations

A reduction in heritability was identified as the main factor reducing prediction accuracy of F1:2 training sets relative to that of F1 training sets, rather than differences in F1 and F1:2 genetic values. Options for increasing the heritability of F1:2 phenotypes include testing in more environments and/or replications and possibly in larger field plots (Rebetzke *et al*. 2014). Alternatively, the lower heritability could partly be compensated by increasing training set size (Lorenz 2013). The increase in resource requirements associated with each option might be justifiable with the expected reduction in complexity and cost of seed production.

This study showed that the ability of F1:2 training sets to predict GCA and hybrid performance on the F1 level is expected to be greater, the lower the degree of divergence between heterotic groups. Most crops for which hybrid breeding is considered promising currently lack clearly defined heterotic patterns (Melchinger and Gumber 1998; Zhao *et al*. 2015; Beukert *et al*. 2017), certainly not to the extend present in maize, where many decades of reciprocal-recurrent selection led to a system of divergent, co-evolving heterotic groups (Duvick *et al*. 2004; Technow *et al*. 2014a; Gerke *et al*. 2015). Thus, for many crop species, for which the lack of an efficient and reliable hybridization system is a major impediment (Longin *et al*. 2012), resorting to F1:2 bulks is particularly promising.

It should be noted that for all self-pollinating crop species, the baseline for comparison is not the “true” F1 hybrids considered in this study, but those produced with the help of hybridization systems. Chemical hybridization agents (CHA), for example, often result in only partial sterility of the female parent, meaning that a certain proportion of the harvested seed is in fact the result of a self pollination. The degree with which this happens is genotype dependent (Adugna *et al*. 2004), leading to a potential confounding of genetic effects and F1 seed purity in field trials. In a CMS system, several cycles of backcrossing are required to develop sterile “A-line” versions of the female inbred lines. In early backcrossing generations, the A-lines still contain significant amounts of donor genome and do not accurately reflect the B-line genotype (Ahmadikhah *et al*. 2015). Neither is necessarily a major concern for production of commercial seed, where CHA applications can be optimized for each commercial hybrid and there is enough time for as many cycles of backcrossing as necessary to remove nearly all residual donor genome. Optimization of CHA applications for hundreds or thousands of experimental hybrids used in a training set or for testcross evaluation, however, is unfeasible, and carrying out three or more cycles of backcrossing before testcrossing would increase the total length of the breeding cycle significantly. Thus, the F1 hybrids used in practice for training set development and testcross evaluation are also not expected to reflect “true” F1 performance without bias.

Even in the “worst-case” scenario however, i.e., without the ability of compensating for the reduced heritability of F1:2 bulks and with strong divergence between heterotic groups, should the prediction accuracy observed here for GCA and hybrid performance of tested and untested individuals be high enough to facilitate genetic gain and identification of superior hybrids. The modest accuracy penalty when using F1:2 bulks might therefore be a reasonable price to pay for the prospect of removing the significant complication and resource requirement of a hybridization system from the breeding process.

## LITERATURE CITED

Adugna, A., G. Nanda, K. Singh, and N. Bains, 2004 A comparison of cytoplasmic and chemically-induced male sterility systems for hybrid seed production in wheat (Triticum aestivum L.). Euphytica 135: 297–304.

Ahmadikhah, A., M. Mirarab, M. H. Pahlevani, and L. Nayyeri-pasand, 2015 Marker-assisted backcrossing to develop an elite cytoplasmic male sterility line in rice. Plant Genome 8.

Albrecht, T., V. Wimmer, H. J. Auinger, M. Erbe, C. Knaak, et al., 2011 Genome based prediction of testcross values in maize. Theor Appl Genet 123: 339–350.

Araus, J. L. and J. E. Cairns, 2014 Field high-throughput phenotyping: the new crop breeding frontier. Trends Plant Sci 19: 52–61.

Bennewitz, J. and T. H. E. Meuwissen, 2010 The distribution of QTL additive and dominance effects in porcine F2 crosses. J Anim Breed Genet 127: 171–179.

Bernardo, R., 1996 Best linear unbiased prediction of the performance of crosses between untested maize inbreds. Crop Sci 36: 872–876.

Beukert, U., Z. Li, G. Liu, Y. Zhao, N. Ramachandra, et al., 2017 Genome-based identification of heterotic patterns in rice. Rice 10: 22.

Bohra, A., U. C. Jha, P. Adhimoolam, D. Bisht, and N. P. Singh, 2016 Cytoplasmic male sterility (CMS) in hybrid breeding in field crops. Plant Cell Rep 35: 967–993.

Bosland, P. W., 2005 Second generation (F2) hybrid cultivars for Jalapeño production. HortScience 40: 1679–1681.

Comstock, R. E. and H. F. Robinson, 1948 The components of genetic variance in populations of biparental progenies and their use in estimating the average degree of dominance. Biometrics 4: 254–266.

Comstock, R. E. and H. F. Robinson, 1952 Estimation of average dominance of genes. In Heterosis, pp. 494–516, Iowa State College, Ames, IA.

Cooper, M., C. D. Messina, D. Podlich, L. R. Totir, A. Baumgarten, et al., 2014 Predicting the future of plant breeding: complementing empirical evaluation with genetic prediction. Crop Pasture Sci 65: 311.

Daetwyler, H. D., R. Pong-Wong, B. Villanueva, and J. A. Woolliams, 2010 The impact of genetic architecture on genome-wide evaluation methods. Genetics 185: 1021–1031.

Duvick, D., 1999 Heterosis: feeding people and protecting natural resources. In The genetics and exploitation of heterosis in crops, edited by J. Coors and S. Pandey, pp. 19–29, CSSA, Madison, WI.

Duvick, D., J. Smith, and M. Cooper, 2004 Long-term selection in a commercial hybrid maize breeding program. In Plant Breeding Reviews, edited by J. Janick, pp. 109–152, John Wiley & Sons, Inc., Hoboken, NJ.

Dwivedi, S. L., A. B. Britt, L. Tripathi, S. Sharma, H. D. Upadhyaya, et al., 2015 Haploids: constraints and opportunities in plant breeding. Biotechnol Adv 33: 812–829.

Endelman, J. B., G. N. Atlin, Y. Beyene, K. Semagn, X. Zhang, et al., 2014 Optimal design of preliminary yield trials with genome-wide markers. Crop Sci 54: 48–59.

Endelman, J. B. and J. L. Jannink, 2012 Shrinkage estimation of the realized relationship matrix. G3 2: 1405–1413.

Falconer, D. S. and T. F. C. Mackay, 1996 Introduction to quantitative genetics. Pearson, fourth edition.

Fischer, S., J. Möhring, H. P. Maurer, H. P. Piepho, E. M. Thiemt, et al., 2009 Impact of genetic divergence on the ratio of variance due to specific vs. general combining ability in winter triticale. Crop Sci 49: 2119–2122.

Fritsche-Neto, R., D. Akdemir, and J. L. Jannink, 2018 Accuracy of genomic selection to predict maize single-crosses obtained through different mating designs. Theor Appl Genet 131: 1153–1162.

Fu, Y., T. J. Wen, Y. I. Ronin, H. D. Chen, L. Guo, et al., 2006 Genetic dissection of intermated recombinant inbred lines using a new genetic map of maize. Genetics 174: 1671–1683.

Gardner, C., 1963 Estimates of genetic parameters in cross-fertilizing plants and their implications in plant breeding. In Statistical Genetics and Plant Breeding, volume 982, pp. 225–251, Comittee on Plant Breeding and Genetics of the Agricultural Board at the North Carolina State College Raleigh, N.C.

Gaynor, R. C., G. Gorjanc, A. R. Bentley, E. S. Ober, P. Howell, et al., 2017 A two-part strategy for using genomic selection to develop inbred lines. Crop Sci 57: 2372–2386.

Gerke, J. P., J. W. Edwards, K. E. Guill, J. Ross-Ibarra, and M. D. McMullen, 2015 The genomic impacts of drift and selection for hybrid performance in maize. Genetics 201: 1201–1211.

Giraud, H., C. Bauland, M. Falque, D. Madur, V. Combes, et al., 2017 Reciprocal genetics: identifying QTLs for general and specific combining abilities in hybrids between multiparental populations from two maize (*Zea mays* L.) heterotic groups. Genetics 207: 1167–1180.

Gorjanc, G., M. Battagin, J. F. Dumasy, R. Antolin, R. C. Gaynor, et al., 2016 Prospects for cost-effective genomic selection via accurate within-family imputation. Crop Sci 57: 216–228.

Habier, D., R. L. Fernando, and D. J. Garrick, 2013 Genomic BLUP decoded: a look into the black box of genomic prediction. Genetics 194: 597–607.

Hallauer, A. R. and S. A. Eberhart, 1970 Reciprocal full-sib selection. Crop Sci 10: 315–316.

Heffner, E. L., A. J. Lorenz, J. L. Jannink, and M. E. Sorrells, 2010 Plant breeding with genomic selection: gain per unit time and cost. Crop sci 50: 1681–1690.

Henderson, C. R., 1973 Sire evaluation and genetic trends. J Anim Sci 1973: 10–41.

Hickey, J. M., S. Dreisigacker, J. Crossa, S. Hearne, R. Babu, et al., 2014 Evaluation of genomic selection training population designs and genotyping strategies in plant breeding programs using simulation. Crop Sci 54: 1476–1488.

Hickey, J. M., G. Gorjanc, R. K. Varshney, and C. Nettelblad, 2015 Imputation of single nucleotide polymorphism genotypes in biparental, backcross, and topcross populations with a hidden Markov model. Crop Sci 55: 1934–1946.

Hill, W. G., M. E. Goddard, and P. M. Visscher, 2008 Data and theory point to mainly additive genetic variance for complex traits. PLoS Genet 4: e1000008.

Holland, J. B., 2001 Epistasis and plant breeding. In Plant Breeding Reviews, edited by J. Janick, pp. 27–92, John Wiley & Sons, Inc., Hoboken, NJ.

Huang, X., S. Yang, J. Gong, Y. Zhao, Q. Feng, et al., 2015 Genomic analysis of hybrid rice varieties reveals numerous superior alleles that contribute to heterosis. Nature Commun 6: 6258.

Jan, H. U., A. Abbadi, S. Lücke, R. A. Nichols, and R. J. Snowdon, 2016 Genomic prediction of testcross performance in canola (Brassica napus). PLOS ONE 11: e0147769.

Janick, J., 1998 Hybrids in horticultural crops. CSSA Special Publication 25: 45–56.

Jannink, J.-L., A. J. Lorenz, and H. Iwata, 2010 Genomic selection in plant breeding: from theory to practice. Brief Funct Genomic Proteomic 9: 166–177.

Jenkins, M. T. and A. M. Brunson, 1932 Methods of testing inbred lines of maize in crossbred combinations. Agron J 24: 523–530.

Kadam, D. C., S. M. Potts, M. O. Bohn, A. E. Lipka, and A. J. Lorenz, 2016 Genomic prediction of single crosses in the early stages of a maize hybrid breeding pipeline. G3 pp. 3442–3453.

Kempe, K. and M. Gils, 2011 Pollination control technologies for hybrid breeding. Mol Breeding 27: 417–437.

Kim, Y.-J. and D. Zhang, 2018 Molecular control of male fertility for crop hybrid breeding. Trends Plant Sci 23: 53–65.

Labate, J., K. R. Lamkey, M. Lee, and W. L. Woodman, 1999 Temporal changes in allele frequencies in two reciprocally selected maize populations. Theor Appl Genet 99: 1166–1178.

Larièpe, A., B. Mangin, S. Jasson, V. Combes, F. Dumas, et al., 2012 The genetic basis of heterosis: multiparental quantitative trait loci mapping reveals contrasted levels of apparent overdominance among traits of agronomical interest in maize (Zea mays L.). Genetics 190: 795–811.

Larièpe, A., L. Moreau, J. Laborde, C. Bauland, S. Mezmouk, et al., 2017 General and specific combining abilities in a maize (Zea mays L.) test-cross hybrid panel: relative importance of population structure and genetic divergence between parents. Theor Appl Genet 130: 403–417.

Longin, C. F. H., X. Mi, and T. Würschum, 2015 Genomic selection in wheat: optimum allocation of test resources and comparison of breeding strategies for line and hybrid breeding. Theor Appl Genet 128: 1297–1306.

Longin, C. F. H., J. Mühleisen, H. P. Maurer, H. Zhang, M. Gowda, et al., 2012 Hybrid breeding in autogamous cereals. Theor Appl Genet 125: 1087–1096.

Lorenz, A. J., 2013 Resource allocation for maximizing prediction accuracy and genetic gain of genomic selection in plant breeding: a simulation experiment. G3 3: 481–491.

Massman, J. M., A. Gordillo, R. E. Lorenzana, and R. Bernardo, 2013 Genomewide predictions from maize single-cross data. Theor Applt Genet 126: 13–22.

Melchinger, A. E., 1999 Genetic diversity and heterosis. In The genetics and exploitation of heterosis in crops, edited by J. Coors and S. Pandey, pp. 99–118, ASA, CSSA, and SSSA, Madison, WI.

Melchinger, A. E. and R. K. Gumber, 1998 Overview of heterosis and heterotic groups in agronomic crops. In Concepts and Breeding of Heterosis in Crop Plant, pp. 29–44, CSSA.

Melchinger, A. E., C. F. H. Longin, H. F. Utz, and J. C. Reif, 2005 Hybrid maize breeding with doubled haploid lines: quantitative genetic and selection theory for optimum allocation of resources. In Proceedings of the forty first annual Illinois corn breeders’ School 2005, pp. 8–21, Urbana-Champaign, Illinois, USA.

Meuwissen, T. H. E., B. J. Hayes, and M. E. Goddard, 2001 Prediction of total genetic value using genome-wide dense marker maps. Genetics 157: 1819–1829.

Mikel, M. A. and J. W. Dudley, 2006 Evolution of North American dent corn from public to proprietary germplasm. Crop Sci 46: 1193–1205.

Mood, A. M., F. A. Graybill, and C. B. Duane, 1973 Introduction to the theory of statistics. McGraw Hill, third edition.

Müller, D., F. Technow, and A. E. Melchinger, 2015 Shrinkage estimation of the genomic relationship matrix can improve genomic estimated breeding values in the training set. Theor Appl Genet 128: 693–703.

Pérez, P. and G. d. l. Campos, 2014 Genome-wide regression & prediction with the BGLR statistical package. Genetics 198: 483– 495.

Poland, J. A. and T. W. Rife, 2012 Genotyping-by-sequencing for plant breeding and genetics. Plant Genome 5: 92–102.

R Core Team, 2018 R: a language and environment for statistical computing. R Foundation for Statistical Computing, Vienna, Austria.

Radoev, M., H. C. Becker, and W. Ecke, 2008 Genetic analysis of heterosis for yield and yield components in rapeseed (Brassica napus L.) by quantitative trait locus mapping. Genetics 179: 1547–1558.

Rebetzke, G. J., R. T. A. Fischer, A. F. v. Herwaarden, D. G. Bonnett, K. Chenu, et al., 2014 Plot size matters: interference from intergenotypic competition in plant phenotyping studies. Functional Plant Biol 41: 107–118.

Reif, J. C., F. M. Gumpert, S. Fischer, and A. E. Melchinger, 2007 Impact of interpopulation divergence on additive and dominance variance in hybrid populations. Genetics 176: 1931–1934.

Riedelsheimer, C., J. B. Endelman, M. Stange, M. E. Sorrells, J. L. Jannink, et al., 2013 Genomic predictability of interconnected biparental maize populations. Genetics 194: 493–503.

Rincent, R., D. Laloe, S. Nicolas, T. Altmann, D. Brunel, et al., 2012 Maximizing the reliability of genomic selection by optimizing the calibration set of reference individuals: comparison of methods in two diverse groups of maize inbreds (Zea mays L.). Genetics 192: 715–728.

Schnell, F., 1965 Die Covarianz zwischen Verwandten in einer gen-orthogonalen Population. Biometrische Zeitung 7: 1–49.

Schön, C. C., B. S. Dhillon, H. F. Utz, and A. E. Melchinger, 2010 High congruency of QTL positions for heterosis of grain yield in three crosses of maize. Theor Appl Genet 120: 321–332.

Schopp, P., D. Müller, F. Technow, and A. E. Melchinger, 2017 Accuracy of genomic prediction in synthetic populations depending on the number of parents, relatedness, and ancestral linkage disequilibrium. Genetics 205: 441–454.

Semel, Y., J. Nissenbaum, N. Menda, M. Zinder, U. Krieger, et al., 2006 Overdominant quantitative trait loci for yield and fitness in tomato. Proc Natl Acad Sci 103: 12981–12986.

Shull, G. H., 1908 The composition of a field of maize. J Hered os-4: 296–301.

Silva Dias, J. C., 2010 Impact of improved vegetable cultivars in overcoming food insecurity. Euphytica 176: 125–136.

Sprague, G. F. and L. A. Tatum, 1942 General vs. specific combining ability in single crosses of corn. Agron J 34: 923–932.

Stuber, C. W., 1970 Estimation of genetic variances using inbred relatives. Crop Sci 10: 129–135.

Stuber, C. W. and C. C. Cockerham, 1966 Gene effects and variances in hybrid populations. Genetics 54: 1279–1286.

Tardieu, F., L. Cabrera-Bosquet, T. Pridmore, and M. Bennett, 2017 Plant phenomics, from sensors to knowledge. Curr Biol 27: 770– 783.

Technow, F., 2013 hypred: Simulation of genomic data in applied genetics. R package version 0.4.

Technow, F. and J. P. Gerke, 2017 Parent-progeny imputation from pooled samples for cost-efficient genotyping in plant breeding. PLOS ONE 12: e0190271.

Technow, F., C. Riedelsheimer, T. A. Schrag, and A. E. Melchinger, 2012 Genomic prediction of hybrid performance in maize with models incorporating dominance and population specific marker effects. Theor Appl Genet 125: 1181–1194.

Technow, F., T. A. Schrag, W. Schipprack, E. Bauer, H. Simianer, et al., 2014a Genome properties and prospects of genomic prediction of hybrid performance in a breeding program of maize. Genetics 197: 1343–1355.

Technow, F., T. A. Schrag, W. Schipprack, and A. E. Melchinger, 2014b Identification of key ancestors of modern germplasm in a breeding program of maize. Theor Appl Genet 127: 2545–2553.

VanRaden, P. M., 2008 Efficient methods to compute genomic predictions. J Dairy Sci 91: 4414–4423.

Whitford, R., D. Fleury, J. C. Reif, M. Garcia, T. Okada, et al., 2013 Hybrid breeding in wheat: technologies to improve hybrid wheat seed production. J Exp Bot 64: 5411–5428.

Windhausen, V. S., G. N. Atlin, J. M. Hickey, J. Crossa, J. L. Jannink, et al., 2013 Effectiveness of genomic prediction of maize hybrid performance in different breeding populations and environments. G3 2: 1427–1436.

Wu, Y. T., J. M. Yin, W. Z. Guo, X. F. Zhu, and T. Z. Zhang, 2004 Heterosis performance of yield and fibre quality in F1 and F2 hybrids in upland cotton. Plant Breeding 123: 285–289.

Würschum, T., J. C. Reif, T. Kraft, G. Janssen, and Y. Zhao, 2013 Genomic selection in sugar beet breeding populations. BMC Genet 14: 85.

Xu, S., 2013 Mapping quantitative trait loci by controlling polygenic background effects. Genetics 195: 1209–1222.

Zhao, Y., Z. Li, G. Liu, Y. Jiang, H. P. Maurer, et al., 2015 Genome-based establishment of a high-yielding heterotic pattern for hybrid wheat breeding. Proc Natl Acad Sci pp. 15624–15629.

