## Supplemental File 2 for "Use of F2 bulks in training sets for genomic prediction of combining ability and hybrid performance"

### Correlation of F1 and F1:2 GCA effects under dominant gene action

The correlation between GCA effects derived from F1 hybrids and F1:2 bulks with a single locus with dominant gene action is expected to be either 1 or -1 in the heterotic group with lower frequency of the trait increasing allele. Values of -1 in particular are expected within a narrow band of degree of dominance by allele frequency divergence combinations (Figure 3B in the main text).

A small numerical example might help to understand the nature of this phenomenon. Single locus genotypes of 10,000 interpopulation hybrids were simulated by drawing 10,000 samples from a Bernoulli distribution with frequency $p_m$, representing the alleles contributed by the parents from the male pool and combining them at random with 10,000 samples from a Bernoulli distribution with frequency parameter $p_f$, representing the female pool. Different values of $p_m$ and $p_f$ were considered to represent different levels of allele divergence. The '1' allele from each group is taken to be the reference allele and will be denoted $B_1$ for consistency of notation with the main text. The alternate allele is $B_2$. The homozygous effect of the locus is $a = 1$ and the heterozygous effect $d = 2.75$. The genotypic values of the F1 hybrids follow directly by assigning values of $a$, $d$ and $-a$ according to the hybrid genotype. The genotypic values of the corresponding F1:2 bulks can be calculated in the same way, except that now $d/2$ has to be used instead of $d$, because 50% of the segregants in the bulks of heterozygous F1 hybrids will have a heterozygous genotype, while the contribution of the 25% segregants with genotypic value of $a$ cancels with the contribution of the 25% with values of $-a$.

Three different divergence levels will be considered, 0.025, 0.70 and 0.95, with $p_f$ receiving the lower allele frequency (e.g., $p_f = 0.15$ and $p_m = 0.85$ at a divergence of 0.70). Only the GCA of the female parents will be considered here, because the described phenomenon affects only the heterotic group with the lower frequency of the trait increasing allele.

With a single, biallelic locus, there will be two distinct GCA values, corresponding to females homozygous for the $B_1$ allele and those homozygous for the $B_2$ allele. Let those effects be denoted as $gca_1$ and $gca_2$. The GCA effects can be calculated as $gca_1 = (1 - p_f)(a + d((1 - p_m) - p_m))$ and $gca_2 = -p_f(a + d((1 - p_m) - p_m))$, or alternatively from the least-squares regression of F1 genotypic values onto the number of $B_1$ alleles contributed by the female parent (either 0 or 1). The slope of this regression line also indicates whether $B_1$ (positive slope) or $B_2$ (negative slope) genotypes have higher GCA effects. For the F1:2 bulks GCA effects can be calculated analogously, except that again $d/2$ has to be used.

**Low divergence** When both heterotic groups have a very similar allele frequency ($p_f = 0.4875$, $p_m = 0.5125$), the distribution in F1 (left) and F1:2 (right) is similar and the GCA values are positively correlated, i.e., $gca_1 > gca_2$ in both (Figure 1).

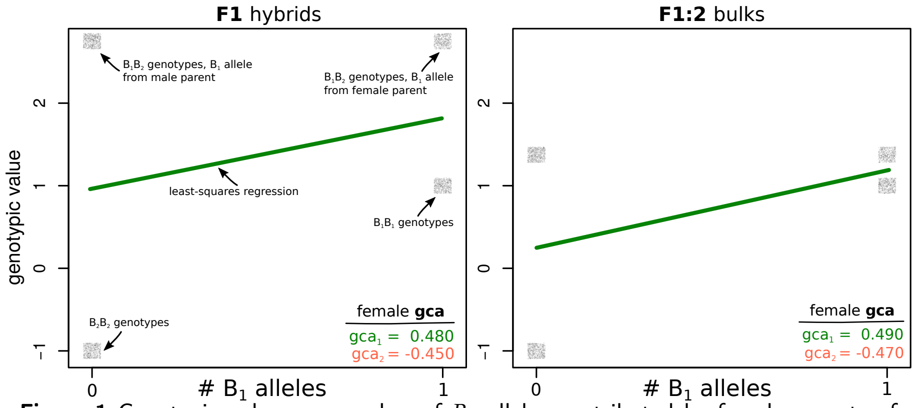


**Figure 1** Genotypic value vs. number of $B_1$ alleles contributed by female parents of hybrids when $p_f = 0.4875$ and $p_m = 0.5125$. The points are 'jittered' along the x and y axes to prevent over-plotting. The colour of the regression line indicates the sign of the slope (green = positive, red = negative).

**High divergence** When both heterotic groups have very divergent allele frequencies ($p_f = 0.025$, $p_m = 0.975$) most hybrids are heterozygous and receive the $B_1$ allele from the males and the $B_2$ from the females (Figure 2). The cluster of heterozygous genotypes in the upper left corner in the figure below thus dominates the distribution and 'pulls' on the slope, making it negative in both F1 and F1:2. The GCA values are again positively correlated because now $gca_1 < gca_2$ in both F1 and F1:2.

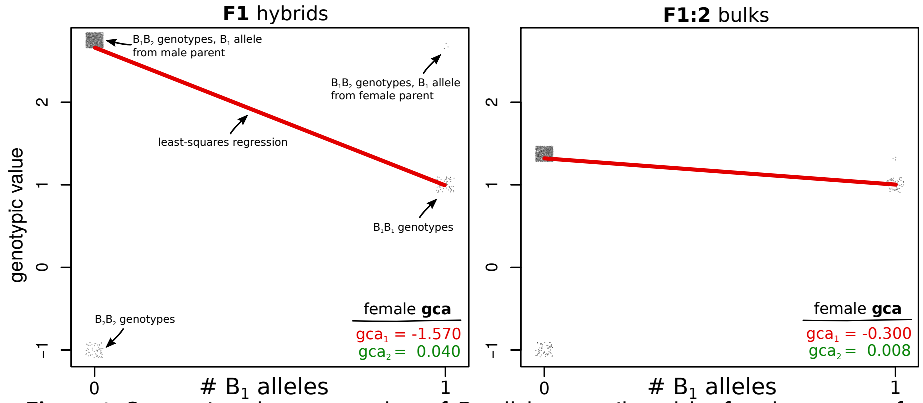


**Figure 2** Genotypic value vs. number of $B_1$ alleles contributed by female parents of hybrids when $p_f = 0.025$ and $p_m = 0.975$. The points are 'jittered' along the x and y axes to prevent over-plotting. The colour of the regression line indicates the sign of the slope (green = positive, red = negative).

**Intermediate divergence** In the intermediate case ($p_f = 0.15$, $p_m = 0.85$), the cluster of heterozygous hybrids that receive the $B_1$ allele from the male parent and the $B_2$ allele from the female parent is still dominant in the F1 and 'pulls' the slope upwards and makes it negative (Figure 3). However, because the weight of this cluster is not as strong as in the previous case (because its frequency is somewhat lower), this 'pull' is not strong enough in the F1:2, where the difference in genotypic value between those hybrids and the hybrids with homozygous genotypes is not as big as in the F1. Thus, the slope remains slightly positive. This results in a negative correlation between F1 and F1:2 GCA, because in the former $gca_1 < gca_2$ while in the latter $gca_1 > gca_2$.

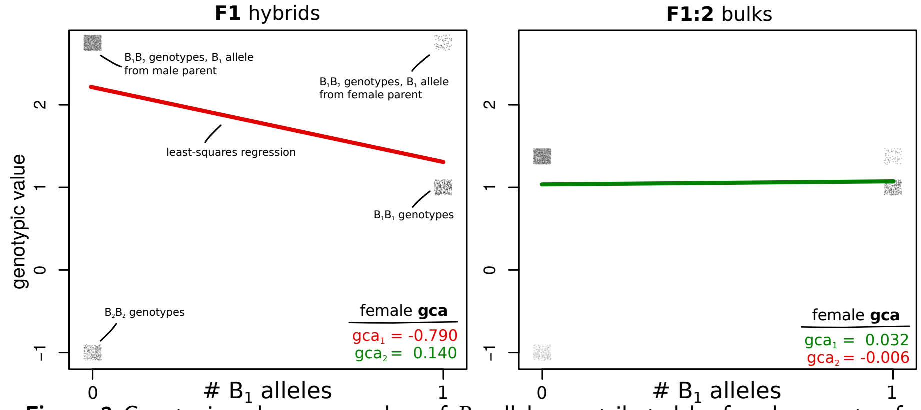


**Figure 3** Genotypic value vs. number of $B_1$ alleles contributed by female parents of hybrids when $p_f = 0.15$ and $p_m = 0.85$. The points are 'jittered' along the x and y axes to prevent over-plotting. The colour of the regression line indicates the sign of the slope (green = positive, red = negative).

This intermediate case corresponds to the 'band' of -1 correlations in Figure 3B. Where it is exactly, depends also the value of $d$. The higher $d$, the sooner will the F1:2 follow the trend of the F1. For example, with $d = 3$, both slopes would already be negative at this intermediate divergence level.
